# Transcriptome and translatome comparison of tissues from *Arabidopsis thaliana*

**DOI:** 10.1101/2023.07.22.550136

**Authors:** Isabel Cristina Vélez-Bermúdez, Wen-Dar Lin, Shu-Jen Chou, Ai-Ping Chen, Wolfgang Schmidt

**Affiliations:** Institute of Plant and Microbial Biology, Academia Sinica, Taipei 11529, Taiwan; Institute of Plant and Microbial Biology, Bioinformatics Core Lab, Academia Sinica, Taipei 11529, Taiwan; Institute of Plant and Microbial Biology, Genomic Technology Core, Academia Sinica, Taipei 11529, Taiwan

## Abstract

Translation is one of the multiple complementary steps that orchestrates gene activity. In contrast to the straightforwardness of transcriptional surveys, genome-wide profiles of the translational landscape of plant cells remain technically challenging and are less well explored. Protein-coding genes are expressed at a variable degree of efficiency, resulting in pronounced discordance among the regulatory levels that govern gene activity. Ribo-Seq provides an extremely useful tool for estimating translation efficiency, but limited data sets are available for plants. Here, we provide comparative inventories of expressed and translated RNA populations, generated by mRNA sequencing (RNA-Seq) and ribosome footprinting (Ribo-Seq) of shoots and roots of *Arabidopsis thaliana* seedlings. Our data set provides information on the translational fitness of protein-coding mRNAs and lncRNAs that may aid in obtaining a comprehensive picture of the regulatory levels governing genes activity across the genome.

## Background & Summary

Gene expression is intricately regulated by transcription, translation, and degradation of the products of these processes, aligning gene activity to the prevailing conditions. Measuring transcription at essentially saturating resolutions has become more attainable and affordable through the advancements in high-throughput technologies such as microarrays and mRNA sequencing (RNA-Seq). Such resolutions are yet a challenge to discovery-based proteomics approaches relying on mass spectrometry, a limitation that is due to the difficulty to detect instable and low-abundance proteins.

Besides incomplete coverage of the proteome, a drawback inherent to proteomic surveys is the fact that the steady-state concentration of proteins is not only determined by the translation rate but heavily affected by their stability^1^, which may lead to erroneous assumptions of the translation rate of a given transcript. The supposition that proteomic profiles are significantly biased by protein degradation is corroborated by the fact that in both yeast and *Arabidopsis* changes in transcript abundance correlated well with alterations in protein levels when upregulated genes were considered, while for downregulated genes no such correlation was observed^1-3^.

Proteins are major players in the orchestration of biological activity, ultimately governing physiological and developmental processes. Translation is energetically costly and, therefore, sophisticatedly regulated to allow for rapid and plastic decision-making. Prioritization of responses to environmental or internal cues defines phenotypic readouts, making such decisions particularly important for plants, which cannot escape unfavorable conditions. Techniques aimed at interrogating the translatome (i.e., the assemblage of mRNAs associated with ribosomes) such as polysome profiling or ribosome footprinting (Ribo-Seq) in combination with highly parallel sequencing were designed to investigate particular regulatory processes that govern translation, indirectly addressing the enigma of the notoriously low concordance of mRNA and protein expression. The gap between transcript and protein abundance is particularly wide in plants, possibly owing to the necessity of highly plastic responses to environmental signals^4^. While in mammalian systems about 40% of gene expression variation has been attributed to transcriptional control, in plants estimates are rather close to 10%^5^, suggesting a large contribution of other factors such as stochastic noise, degradation, and translational regulation. Thus, investigating the mechanisms that control protein abundance to understand plant function and development appears to be an obligatory and promising approach.

Interrogations into the *in vivo* translational landscape of plants enable the discovery of regulatory mechanisms of translational control, the identification of novel translated short open reading frames (sORFs), previously undiscovered upstream open reading frames (uORFs), non-AUG start codons, the determination of the frame and length of the translated regions, and, consequently, the discovery of novel proteins. Monitoring translation is, however, technically more challenging than measurements of the transcriptome. The invention of the Ribo-Seq methodology, originally developed in the yeast *Saccharomyces cerevisiae*^6^, was a big leap forward towards the understanding of translational regulation, but data on what is when and how efficiently translated are still scarce, in particular in plants^7^. The incomplete coverage of proteomic approaches renders investigations on translational control of transcripts difficult, even when label-free techniques are employed that allow estimates of absolute protein concentrations. Ribo-Seq provides insights into the density and precise location of translating ribosomes over the entire length of the transcripts, allowing massively parallel sequencing of ribosome-protected footprints (RPFs) after subjecting the remainder of the mRNAs to nucleolytic digestion. Ribo-Seq not only grants access to the inventory of the population of transcripts bounded by ribosomes and thus estimates the ‘protein potential’ of the cell, but also provides a snapshot of the precise position of the ribosomes at the time when this process (artificially) came to a halt.

The aim of the present study was to provide a comprehensive inventory of Ribo-Seq-generated RPFs form roots and shoots of the reference plant *Arabidopsis thaliana*.

The abundance of RPFs is normalized to the steady-state level of mRNA abundance estimated by RNA-Seg, allowing estimates on the translational fitness of transcripts derived from protein-coding genes. The experimental setup is depicted in Figure 1. Plants were grown on sterile, agar-solidified media for 14 days, dissected into shoots and roots, and immediately frozen in liquid nitrogen. After cell lysis, RNA-Seq samples were generated by extracting total RNA and library construction and subjected to next-generation sequencing. For Ribo-Seq, the samples were digested with RNAse I followed by sucrose cushion ultracentrifugation, size selection, ribosome footprint purification, library construction, and sequencing. In a final step, both data sets were analysed and compared to estimate the translation efficiency of the expressed transcripts.

**Fig. 1.**
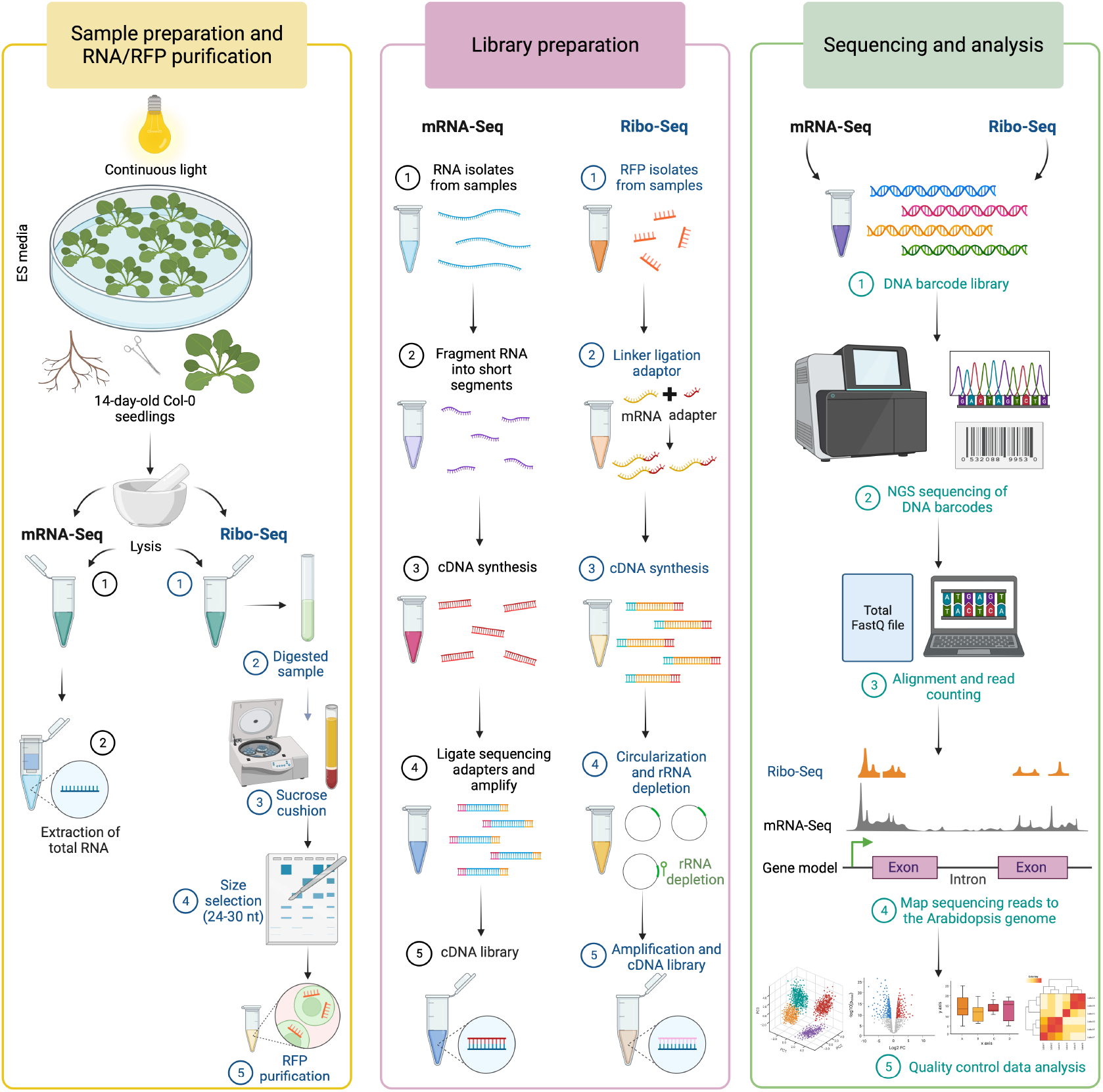
Overview of the experimental design. The principal steps of the RNA-Seq and Ribo-Seq analyses are depicted in order of their execution. Protocols for Ribo-Seq and RNA-Seq are detailed in the Methods section.

## Methods

### Plant material, growth conditions, and sample collection

Seeds of the Col-0 accession of *Arabidopsis thaliana* were surface sterilized by soaking them in 35% sodium hypochlorite for 5 min, followed by five rinses in sterile water (5 minutes each). The seeds were then placed in a growth medium composed of 5 mM KNO3 (7757-79-1, Merck), 2 mM MgSO_4_ × 7H_2_O (10034-99-8, SIGMA), 2 mM Ca(NO_3_)_2_ ×_2_4H_2_O (13477-34-4, Sigma), 2.5 mM KH_2_PO_4_ (7778-77-0, Merck), 70 μM H_3_BO_3_ (B0252, Sigma), 14 μM MnCl_2_ (13446-34-9, Merck), 1 μM ZnSO_4_ (7446-20-0, Merck), 0.5 μM CuSO_4_ (7758-98-7, Sigma), 0.01 μM CoCl_2_ (7646-79-9, Sigma), 0.2 μM Na_2_MoO_4_ (10102-40-6, Merck); and 40 μM ethylenediaminetetraacetic acid iron(III) sodium salt (NaFe-EDTA; EDFS, Sigma), solidified with 0.4% Gelrite Pure (Supermacy Instrument CO., LTD). Sucrose (1.5% (w/v); Sigma) and 1 g/L MES (M8250, Sigma) were added, and the pH was adjusted to 5.5 with KOH (1310-58-3, Merck). The plates containing the seeds were stratified for 2 days at 4°C in the dark, transferred to a growth chamber, and grown at 22°C under continuous illumination (50 mmol m^-2^ s^-2^; Philips TL lamps) and 70% relative humidity. After 14 days, samples were collected for the experiments. The seedlings were gently removed from the plates, shoots and roots were separated using a razor blade, and the tissues were flash-frozen in liquid nitrogen. Samples were kept at −80°C until RNA or RPF extraction. For RNA-Seq, twelve shoots and twenty-five independent roots were bulked to form one biological replicate. For Ribo-Seq, twenty-five shoots and fifty independent roots were bulked for each sample. Purified mRNA purification and ribosome footprints were collected in biological duplicates.

### RNA-Seq

Stranded RNA-Seq analysis was conducted in duplicates. Total RNA was extracted from 100 mg of roots or shoots using the RNeasy Plant Mini Kit (Qiagen) following the supplier’s instructions. Paired-end cDNA libraries were constructed from 4 μg of total RNA from root and shoot samples, with insert sizes ranging from 100 to 600 bp using a commercial kit (TruSeq stranded mRNA Sample Preparation Kit; Illumina Inc., San Diego, CA, USA). The libraries were sequenced for 100 bases using an Illumina Hiseq 2500 SE100 system.

### Ribo-Seq

Ribosome footprints and library construction were performed as described in a previous study^8^ in duplicates with modifications. Briefly, 200 mg of *Arabidopsis* roots or shoot were resuspended in 600 μL of lysis buffer (polysome buffer; 20 mM Tris-HCl pH 7.4, 150 mM NaCl, 5 mM MgCl_2_, 1 mM DTT supplemented with 100 μg/ml cycloheximide, 1% Triton X-100, and Turbo DNase I 25 U/mL), incubated for 15 min on ice with vortex every five minutes, and centrifuged at 16,000 × g for 15 min at 4°C. Then, 500 μL of lysate were digested with 12.5 μL RNase I (100 U/μL) for 45 min at room temperature with gentle mixing on a nutator. Subsequently, 16.66 μL of SUPERase*In RNase Inhibitor were added to stop the nuclease digestion. The ribosome pellet was recovery by centrifugation at 70,000 rpm at 4°C for 4 hours in a TLA 100.4 rotor, and the ribosomal footprints were purified using TRIzol® reagent according the manufacturer’s instructions. RFPs were then separated on a 15% urea denaturing-PAGE gel, and gel slices corresponding to 24–30 nts were excised. RFP-derived RNAs were eluted and precipitated followed by library construction. Four biotinylated oligonucleotides, Oligo 1: 5’/5BioTEG/CATAAACGATGCCGACCAGGGATCAGCGG-3’, Oligo 2: 5’ /5BiotinTEG/CTC TGATGATTCATGATAACTCGACGGATCGCATGG-3’, Oligo 3: 5’ /5BiotinTEG/CATTAGCATGGGATAACATCAT-3’, and Oligo 4: 5’-/5BiotinTEG/TGCCAAGGATGTTTTCATTAATCAAGAACG-3’ were used to remove contaminating rRNA fragments from the libraries. The libraries were sequenced for 50 bases using an Illumina Hiseq 2500 SR50 system.

### Normalization of expression and translational levels

Expression (RNA-Seq) and translational levels (Ribo-Seq) were normalized as RPKM (reads per kilobase of transcript per million reads mapped) using read counts and feature lengths.

### Mapping of RNA-Seq and Ribo-Seq reads

RNA-Seq reads were mapped to the TAIR10 genome using BLAT^9^ with default parameters. RPKM values were computed using the inhouse RackJ toolkit (http://rackj.sourceforge.net/)based on reads mapped with at least 95% identity. Ribo-Seq reads were firstly processed by Cutadapt^10^ for adaptor removal. Cleaned reads were filtered by mapping them to Araport11^11^ non-coding RNAs, where reads with at least 95% identity were removed. The remaining reads were mapped to the Araport11 genome annotation using Bowtie2^12^ accepting alignments with identity at least 95% identity. Unaccepted reads were mapped to junction-spanning sequences using Bowtie2. Junctions were inferred based on RNA-Seq data, each confirmed by a minimum of 10 reads with at least 4 different mapping start positions. Thereby, alignments of Ribo-Seq reads spanning junctions unknown to the Araport11 database were recovered. The remaining reads were directly mapped to the Araport11 genome using BLAT. Alignments with an identity of less than 95% and shorter than 8 bps were removed. All accepted alignments were transferred to the same coordinate of the Araport11 genome.

Multi-reads with two or more best alignments were collected for further filtering. Covering regions were identified and corresponding multi-read counts were computed. As a result, five regions covered by more than 95% of multi-reads were identified and reads with alignments in these regions were removed from the computation. The alignments resulting from this procedure were used for RPKM computation.

### Principal Component Analysis (PCA) analysis

The PCA plot was generated using the R software package.

### Pearson correlation calculations

Pearson correlation analysis was based on log-RPKM and performed for every two biological replicates for roots and shoots (RNA-Seq and Ribo-Seq) using an inhouse Perl script.

### Hierarchical clustering

Heatmaps and hierarchical clustering of detected genes derived from Ribo-Seq and RNA-Seq were generated using Next-Generation Clustered Heat Map Viewer^13^.

### Calculation of length distribution and nucleotide periodicity of RPFs

RPF length distribution and 3-nt periodicity were determined using an inhouse Perl script and Excel. 3-nt periodicity was estimated by counting the base on the 13th bp positions of the Ribo-Seq reads.

### Identification of uORFs

uORFs longer than 10 aa were identified by an AUG start codon. The identified lncRNAs were combined to identify uORFs across the genome.

### Calculation of translation efficiency (TE)

For TE analysis, we considered genes to be expressed with RPKM > 0. TE computation was made using entire transcripts, unique read counts of RNA-Seq and Ribo-Seq, and based on RPKM. For the TE scatter plot in log-log scale, genes with RNA-Seq and/or Ribo-Seq RPKM = 0 in the two biological replicates were excluded. The TE GO enrichment analysis was performed using the ArgiGO v2.0 toolkit web-server^14^. Significantly enriched GO terms were summarized and visualized using REVIGO^15^ as described in a previous study^16^.

### Comparison of the number of genes detected by mass spectrometry, Ribo-seq, and RNA-seq

The comparison among the total detected genes (RPKM > 0) in roots and shoots in the RNA-Seq, Ribo-Seq, and the tier1-canonical proteins^17^ data sets was made using a Venn diagram online tool (https://bioinformatics.psb.ugent.be/webtools/Venn/)^18^.

### Data availability

Raw read FASTQ files, trimmed read FASTQ files, mapped read BAM files (mapped to Araport11), and gene expression in RPKM in Excel files of all samples were deposited at the National Center for Biotechnology Information Sequence Read Archive (NCBI SRA) under study BioProject accession PRJNA990964.

### Technical Validation

#### Experimental design

*A. thaliana* (Col-0) seedlings were growth under controlled conditions for 14 days. In order to perform RNA-Seq and Ribo-Seq experiments, we separately collected shoot and root samples. The plant material was collected without wounding or damaging the plant tissue; undamaged tissue is crucial to obtaining high quality data in particular for Ribo-Seq surveys. For RNA-Seq experiments, we bulked twelve shoots and twenty-five roots, Ribo-Seq necessitates twenty-five shoots and fifty independent roots per replicate, all samples were arranged in duplicates. An overview of the experimental design and explanation of RNA-Seq and Ribo-Seq steps is shown in Figure 1.

### Quality control of the RNA-Seq and Ribo-Seq data

RNA-Seq and Ribo-Seq libraries were generated from roots and shoots in duplicates, and sequenced using Illumina platforms (Table 1). RNA-Seq libraries yielded 44,798,444 (Roots R1) and 45,622,719 reads (Roots R2) from mRNA extracted from roots and 44,650,350 (Shoots R1) and 42,707,942 (Shoots R2) reads from mRNA extracted from shoots. For Ribo-Seq, the number of yielded reads were 58,082,938 (Roots R1), 67,623,514 (Roots R2), 59,738,467 (Shoots R1), and 53,278,422 (Shoots R2). For RNA-Seq samples, reads that mapped to a unique gene were 43,133,955 (Roots R1), 43,716,472 (Roots R2), 43,070,070 (Shoots R1), and 41,126,032 (Shoots R2) with an average length of the trimmed reads between 100 and 101 nt. For Ribo-Seq, reads were considered that directly mapped to more than 95% to the Araport11 genome annotation and those with alignments shorter than 8 bp were removed. For protein-coding genes, the latter step contributed to no more than 1% unique Ribo-Seq reads. For Ribo-Seq, the reads mapped to unique genes were 4,684,123 (Roots R1), 3,965,582 (Roots R2), 5,795,674 (Shoots R1), and 4,310,899 (Shoots R2). It thus appears that the Ribo-Seq library exhibits sufficient reads mapping to annotated CDS, with average length of filtered read between 28-34 nt. However, due to the shorter length of the reads, the number of uniquely mapped reads was about 10-fold lower for Ribo-Seq-derived reads when compared to RNA-Seq reads (Table 1).

**Table 1.**
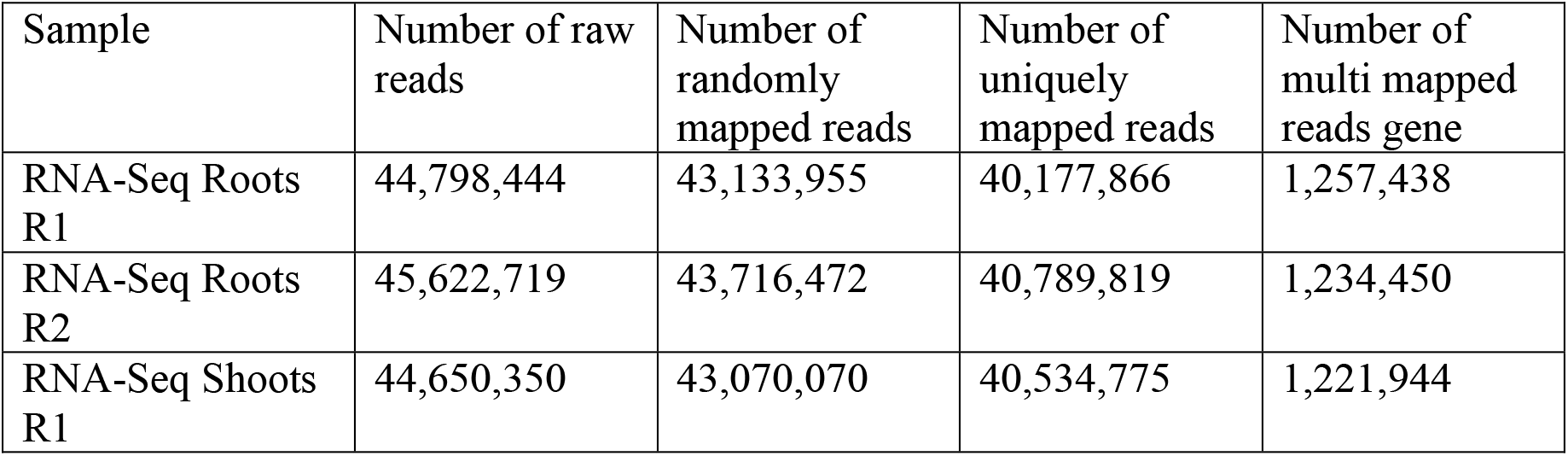
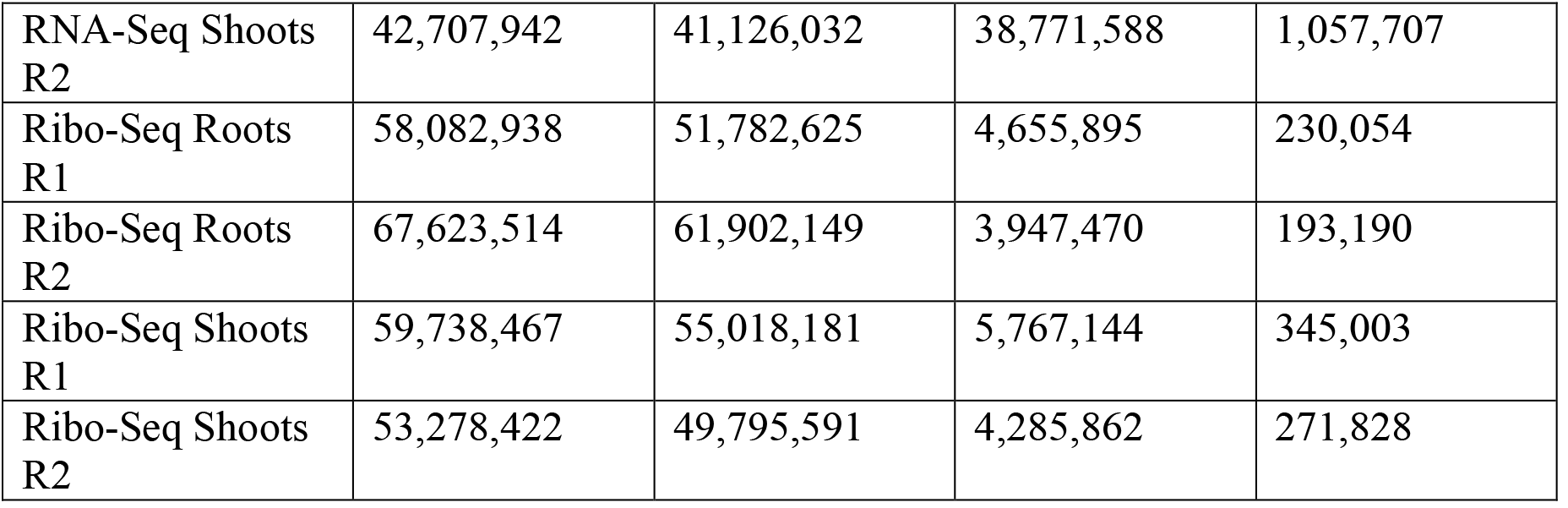
Read numbers of the RNA-Seq and Ribo-Seq data sets.

### Reproducibility of the RNA-Seq and Ribo-Seq data

We performed a Principal Component Analysis (PCA) for the data derived from the RNA-Seq (total mRNA) and Ribo-Seq (RPFs) profiling experiments, using individual biological replicates for each tissue under study (Fig. 2a). The PCA plots shows low bias associated with the biological replicates for both RNA-Seq and Ribo-Seq, supporting the robustness of the data. The results reveal that the variability of the Ribo-Seq replicates was somewhat higher when compared to the high reproducibility of the RNA-Seq data, an observation that is possibly associated with the more complex protocol of the former methodology.

**Fig. 2.**
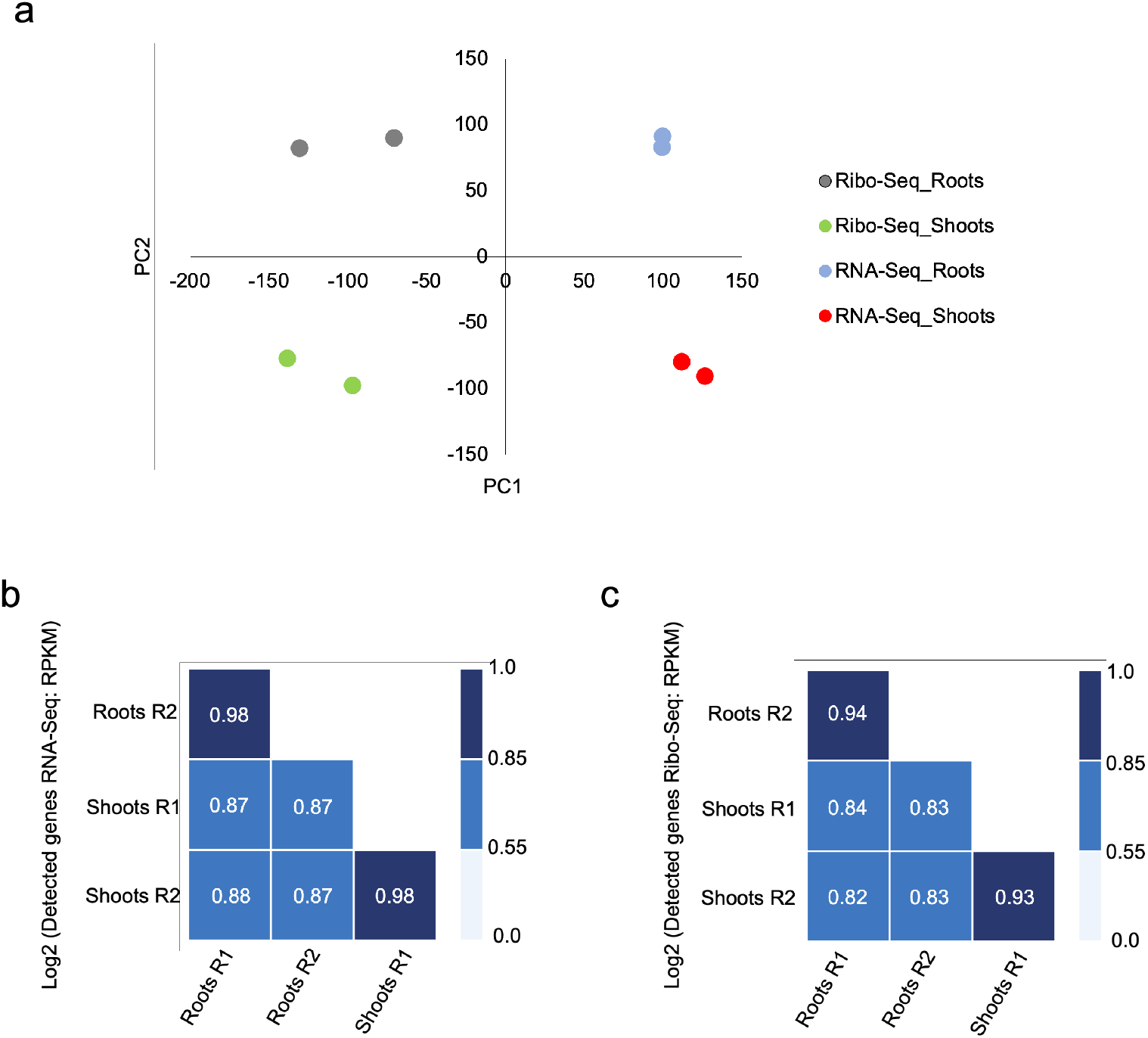
Technical validation. (a) Principal-component analysis (PCA) of the RNA-Seq and Ribo-Seq data derived from roots and shoots. The analysis demonstrates the high repeatability of both surveys. The samples were categorized by the individual biological replicates of each tissue for each technique. (b-c) Pearson correlation coefficient values of the RNA-Seq samples (b) and Ribo-Seq samples (c).

Normalized reads derived from the RNA-Seq and Ribo-Seq surveys were used to define detected genes in the transcriptome and translatome. The reproducibility of both experiments was confirmed by high Pearson correlation between sample pairs (Fig. 2b-c) and hierarchical clustering of the data, which corroborated the robustness of both methods (Fig. 3a-b). Also here, RNA-Seq data appeared to be less noisy and more heterogenic among the biological replicates relative to RNA-Seq data. To further evaluate the quality of the Ribo-Seq data, we analysed the 3-nucleotide (nt) periodicity—a phenomenon that allows predicting open reading frames—using individual biological replicates for roots and shoots (Fig. 4a). The triplet periodicity is one of the most important features of Ribo-Seq data derived from protein-coding genes. Such periodicity is usually not observed in non-coding regions. We computed and aggregated read depths of the first 50 bps from all representative coding models, where read depths were computed based on the 13th base pair of every mapped Ribo-Seq read. In each sample, the plots show a pronounced 3-nt periodicity (Fig. 4a). In addition, the first codon showed stronger signals than the other codons, a further feature that is typically observed in Ribo-Seq reads.

**Fig. 3.**
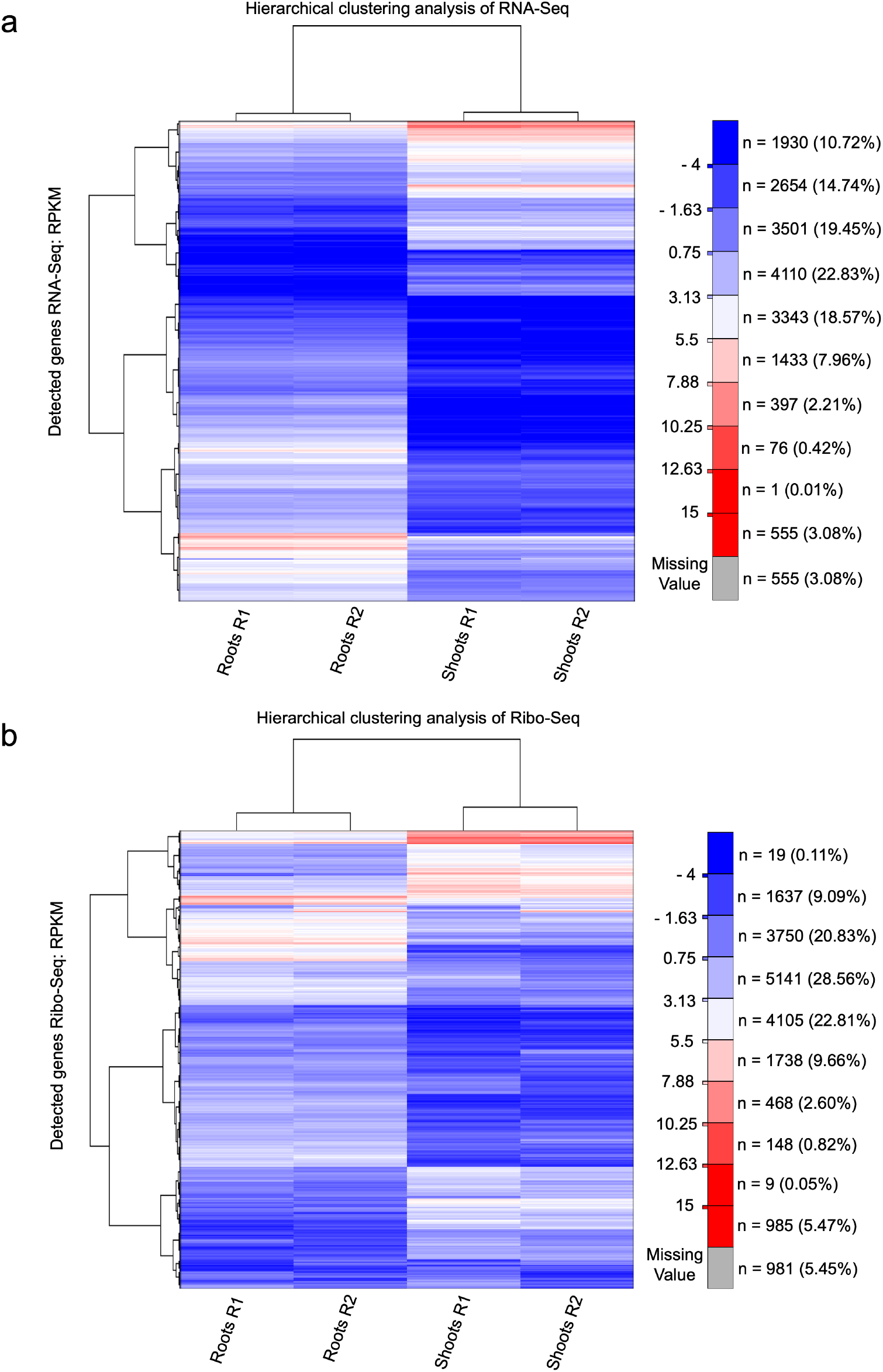
Quality validation of the data. (a) Hierarchical clustering analysis of RNA-Seq samples from roots and shoots. (b) Hierarchical clustering analysis of Ribo-Seq samples from roots and shoots.

**Fig. 4.**
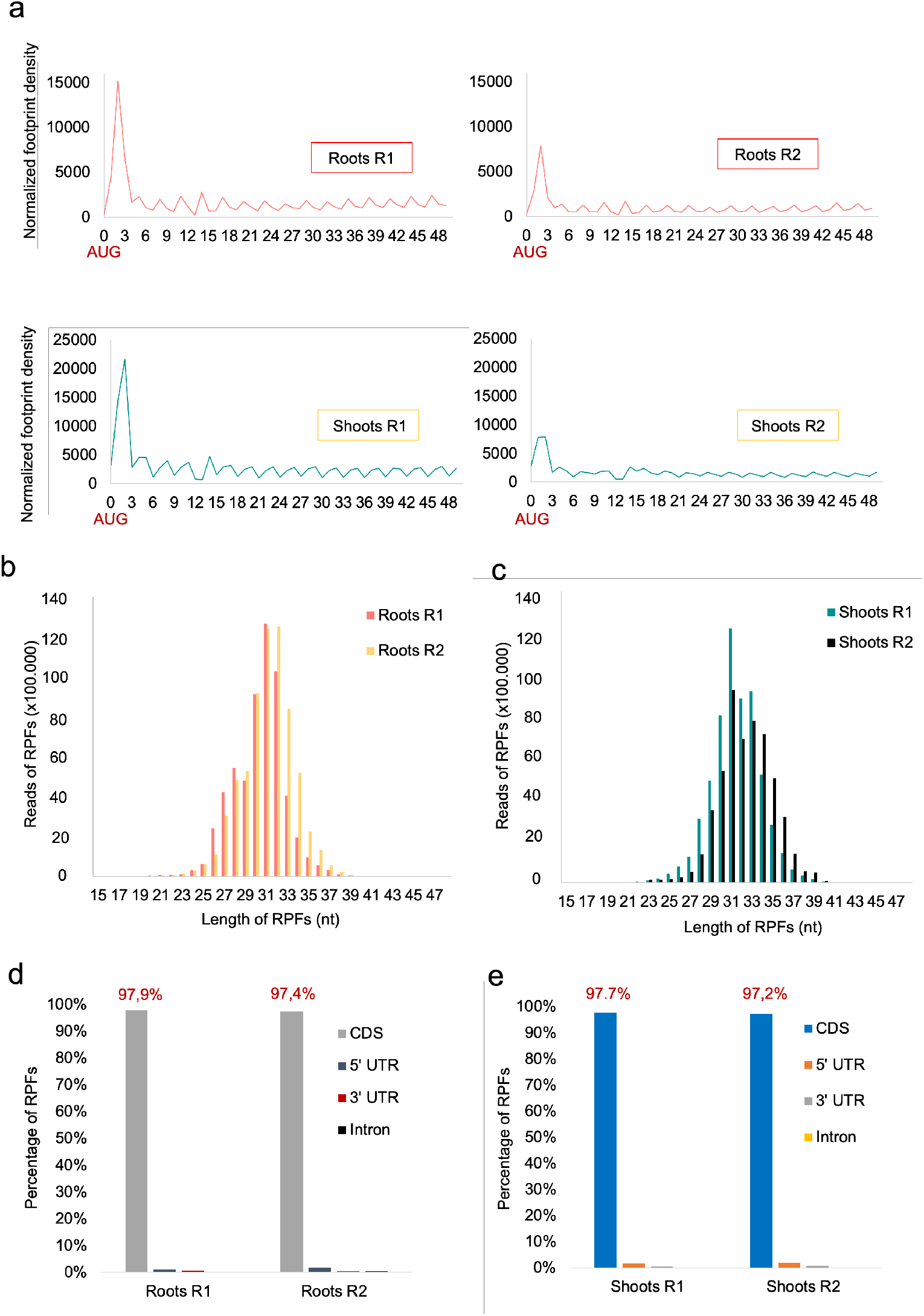
Reproducibility of the data. (a) Three-nucleotide periodicity of the Ribo-Seq data. The position of the ribosomes along the transcripts was computed based on the 13th base pare for individual biological replicates of roots (upper panel) and shoots (lower panel). (b-c) Length distribution of RPFs from roots (b) and shoots (c) showing a peak at 30 nt. (d-e) Percentage of RPFs mapped to CDS, 5’ UTR, 3’ UTR, and introns in samples from roots (d) and shoots (e).

The length of the distribution of RPFs is another feature that we measured to determinate quality of the libraries. We found that the vast majority of RFPs from each biological replicate showed a read length distribution ranging from about 26 nt to 34 nt with a peak at 30 (Fig. 4b-c), a distribution that has been observed in other studies using Arabidopsis tissue^19^. When the RFPs were mapped to the Arabidopsis genome (Fig. 4d-e), 97.9% and 97.4% of the reads for the root samples R1 and R2 localised to annotated CDS, the remaining RFPs were located in the 5’ UTR, 3’ UTR, or in intronic regions. For shoot samples, 97.7% and 97.2% of the reads were located in the CDS while the remainder of RFPs were aligned to 5’ UTR, 3’ UTR, or intronic regions.

### Usage Notes

#### Identification of uORFs

Upstream open reading frames are located in the 5’ UTR regions and thought to regulate protein synthesis by acting as translational repressors. Considering genes defined as ‘detected’ in the Ribo-Seq data set, we estimate that the percentage of uORFs was 26.7% and 29.1% for roots and shoots, respectively. The frequency distribution of uORF lengths was showing similar pattern in roots and shoots (Fig. 5a-b). In total, 112 lncRNAs were detected in roots and shoots. The frequency distribution of uORFs lengths within lncRNAs is shown in Figure 5c-d. Our data indicate that in roots 77.7% (87) of all lncRNAs with one or more uORF are translated, while the number of putatively translated lncRNAs is somewhat lower in shoots (52.7%; 59).

**Fig. 5.**
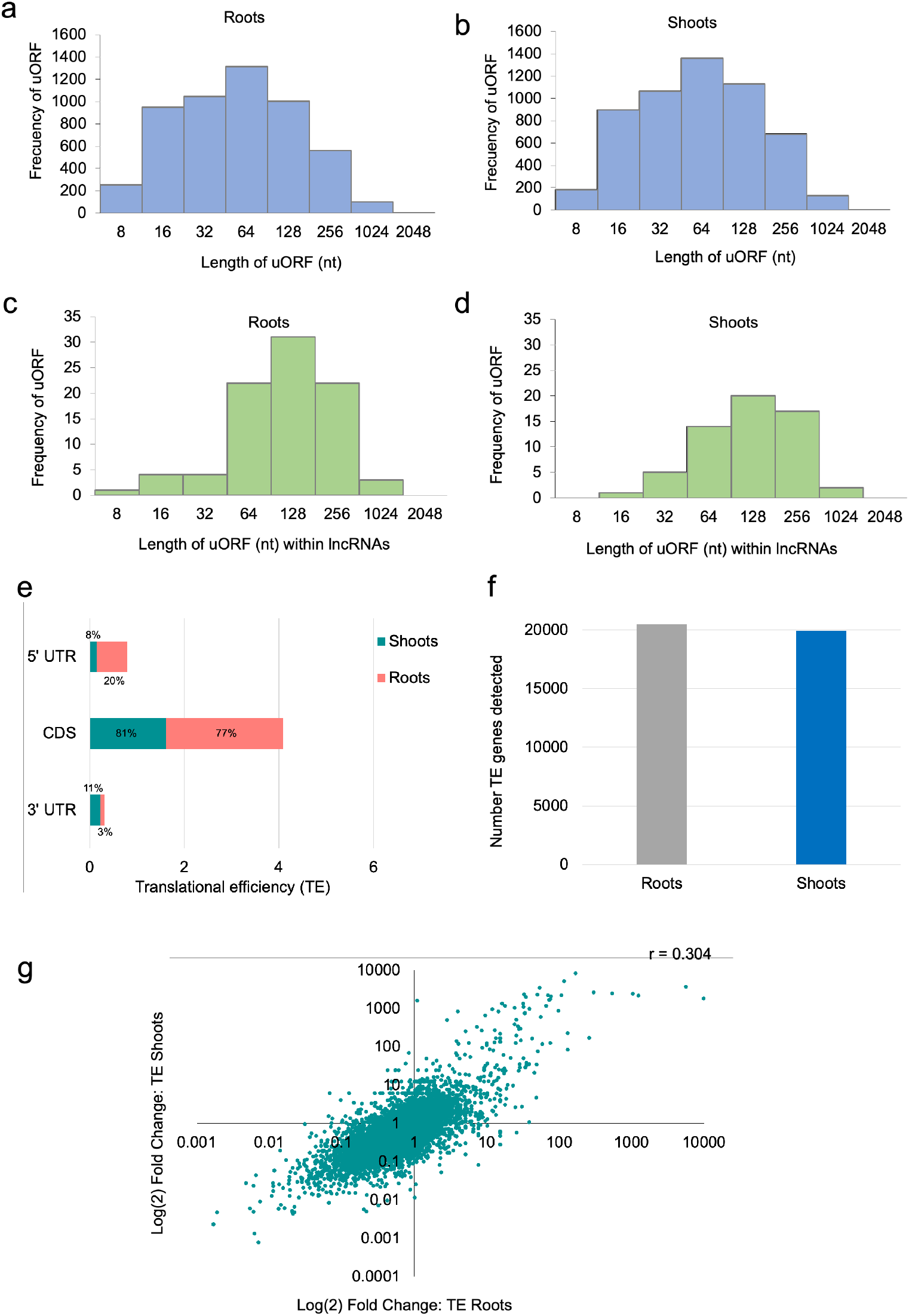
Usage examples. (a-d) Frequency of the length distribution of the translated uORFs in roots (a), shoots (b), within lncRNAs in roots (c), and within lncRNAs in shoots (d). (e) Distribution of the translation efficiency in the 5’ UTR, CDS and 3’ UTR in roots and shoots. (f) Number of TE detected genes in different tissues. (g) Translation efficiency scart plot in log-log scale for roots and shoots.

#### Translation efficiency

Ribo-Seq normalised to RNA-Seq data have been used to study the dynamics of translation efficiency of a group of genes under particular conditions or at a genome wide scale.^7^ We analysed the translation efficiency (TE; ratio of the RPFs over mRNA counts) of genes from roots and shoots in the 5’ UTR, CDS, and 3’ UTR. As expected, highest TE was observed in the CDS with comparable values for shoots (81%) and roots (77%). TE in 5’ UTR and 3’ UTR ranged around 10%, but the 5’UTR of roots displayed a markedly higher ribosome occupancy (Fig. 5e). In both organs, TE could be calculated for an approximately similar number of genes (Fig. 5f).

A log-log scale scatter plot of the combined TEs values from shoots and roots revealed a moderate positive correlation between both organs (Fig. 5g), suggesting that the TE of a particular gene is only weakly dependent on its location. Gene ontology enrichment (biological process) for the genes with the highest TE (top 5% TE) detected in roots and shoots revealed that genes that are efficiently translated are involved in crucial biological process. In both roots and shoots, highest enrichment was observed for genes involved in energy-generating processes. In the case of roots, the categories ‘response to abiotic stimulus’ and ‘transmembrane transport’ were also enriched, indicating processes that are particularly prioritised in roots (Fig. 6a). In shoots, a large number of genes with high TE are involved in nitrogen and phosphorous metabolism. Genes associated with purine metabolism were enriched in both organs (Fig. 6a,b).

**Fig. 6.**
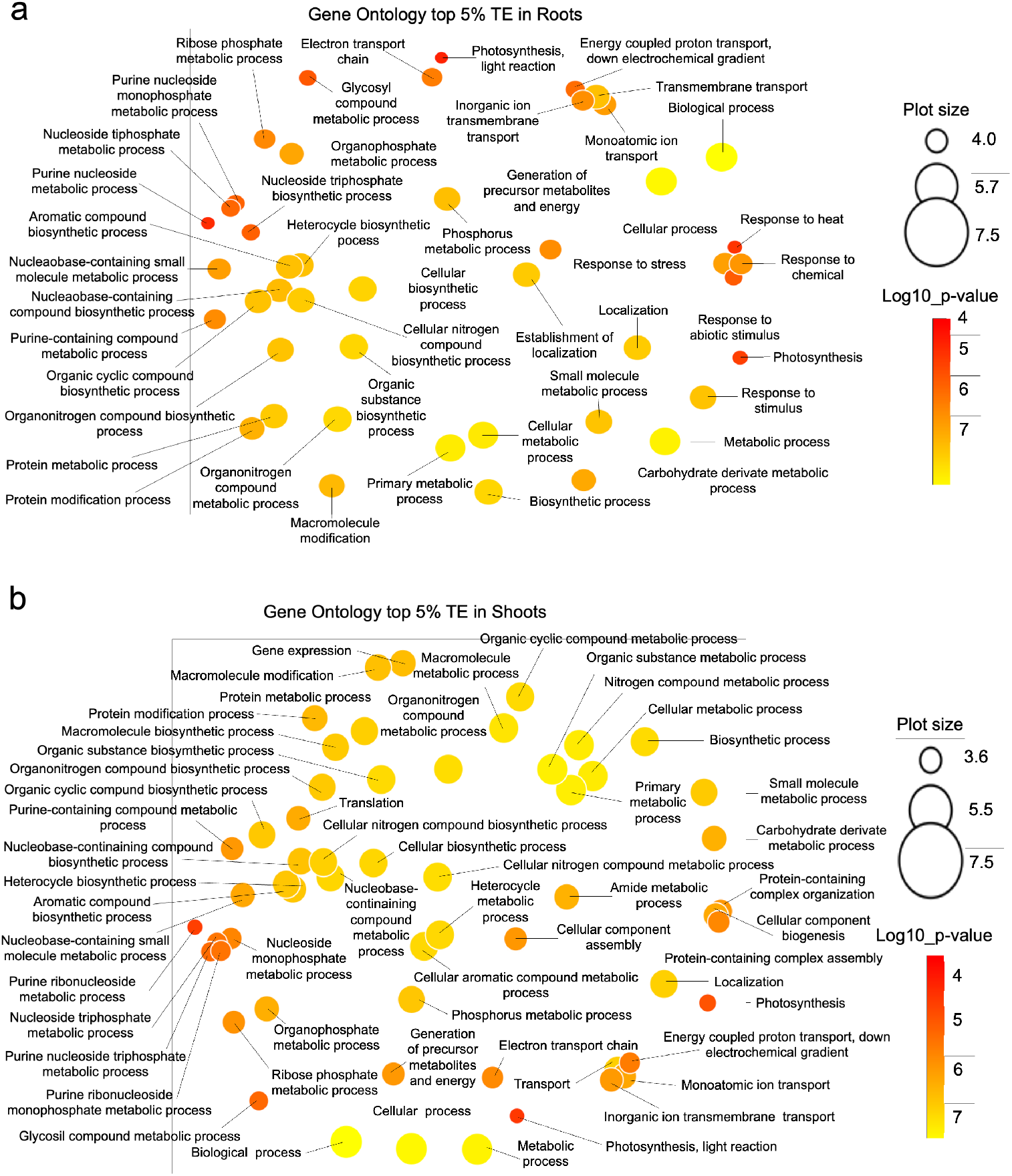
Gene ontology (GO) enrichment within detected genes with the highest (top 5%) TE. (a-b) GO enrichment (biological process) in roots (a) and shoots (b).

#### Robustness of the Ribo-Seq data

A multi-omics comparation using the total list of detected genes (roots and shoots) in the RNA-Seq and Ribo-Seq data sets against a previously published inventory of the Arabidopsis proteome^17^ that comprised proteins with at least two uniquely mapping non-nested peptides showed that a large number (15,734) of genes were found in all three data sets (Fig. 7). A subset of 6,690 genes detected by RNA-Seq was not covered by the proteomic data set. For Ribo-Seq, a subset of 3,847 genes was not comprised in the expressed proteome. A subset of 3,784 genes was comprised in the RNA-Seq and Ribo-Seq data but not in the proteome, suggesting that these genes are translated to low abundant or unstable proteins.

**Fig. 7.**
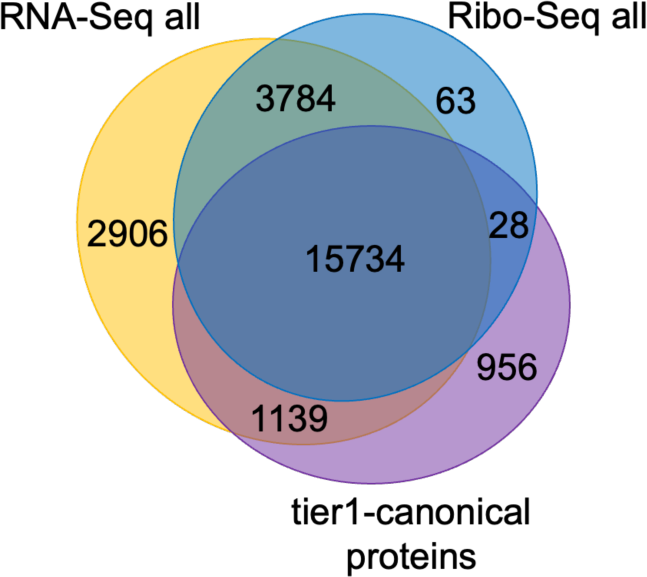
Comparison of the genes detected by RNA-Seq and Ribo-Seq data with expressed proteins. The proteomic data set was derived from a previous study^17^.

## Code availability

All bioinformatics tools used in this work are described in the Methods section.

## Acknowledgements

This work was supported by grants from Academia Sinica (AS-TP-109-L01) and the National Science and Technology Council (111-2313-B-001-01). Figure 1 was created with BioRender.com.

## Author contributions

W.S. and I.C.V-B. designed and conceived the experiments. I.C.V-B., S-J.C. and A-P.C. conducted the experiments. W-D.L. performed the computational analysis. W.S., I.C.V-B. and W-D.L., S-J.C. and A-P.C. analysed the results. I.C.V-B. and W.S. wrote the manuscript.

